# Parkinson’s disease determinants, prediction and gene-environment interactions in the UK Biobank

**DOI:** 10.1101/2020.02.15.950733

**Authors:** Benjamin M. Jacobs, Daniel Belete, Jonathan P Bestwick, Cornelis Blauwendraat, Sara Bandres-Ciga, Karl Heilbron, Ruth Dobson, Mike A. Nalls, Andrew B. Singleton, John Hardy, Gavin Giovannoni, Andrew J. Lees, Anette Schrag, Alastair J Noyce, for The International Parkinson’s Disease Genomics Consortium (IPDGC)

## Abstract

**Objective:** To systematically investigate the association of environmental risk factors and prodromal features with incident Parkinson’s disease (PD) diagnosis and the interaction of genetic risk with these factors. To evaluate existing risk prediction algorithms and the impact of including addition genetic risk on the performance of prediction.

**Methods:** We identified individuals with incident PD diagnoses (n=1276) and unmatched controls (n=500,406) in UK Biobank. We determined the association of risk factors with incident PD using adjusted logistic regression models. A polygenic risk score (PRS) was constructed and used to examine gene-environment interactions. The PRS was also incorporated into a previously-developed prediction algorithm for finding incident cases.

**Results:** Strong evidence of association (P_corr_<0.05) was found between PD and a positive family history of PD, a positive family history of dementia, non-smoking, low alcohol consumption, depression, and daytime somnolence, and novel associations with epilepsy and earlier menarche. Individuals with the highest 10% of PRS scores had increased risk of PD (OR=3.30, 95% CI 2.57-4.24) compared to the lowest risk decile. Higher PRS scores were associated with earlier age at PD diagnosis and inclusion of the PRS in the PREDICT-PD algorithm improved model performance (Nagelkerke pseudo-R^2^ 0.0053, p=6.87×10^−14^). We found evidence of interaction between the PRS and diabetes.

**Interpretation:** Here we used UK Biobank data to reproduce several well-known associations with PD, to demonstrate the validity and predictive power of a polygenic risk score, and to demonstrate a novel gene-environment interaction, whereby the effect of diabetes on PD risk appears to depend on prior genetic risk for PD.

## Introduction

Parkinson’s disease (PD) is the second most prevalent neurodegenerative disorder worldwide and potentially the fastest growing^1^. By the time an individual is diagnosed with PD, a substantial proportion of nigrostriatal neurons have already been lost^2^. There is an urgent clinical and societal need for effective treatments or strategies which will prevent, halt or reverse the progression of PD. Prediction of those at-risk and/or detection at the earliest stages likely represent the best approaches to address this need.

Major progress has been made in understanding the genetic architecture of PD. As for many complex diseases, this began with linkage studies of rare, familial forms of PD which revealed pathogenic roles for *SNCA, PARK2* and *PINK1* genes, and later *LRRK2*, and *GBA*^*3*^. Over the last decade, large genome-wide association studies (GWAS) of sporadic PD have extended our understanding of the genetic architecture of PD to include 90 independent signals, which collectively explain ∼22% of overall PD liability^4^.

Separately, there exists good epidemiological evidence to support a role for potentially-modifiable exposures including pesticide exposure, head injury, and potentially protective factors such as smoking, and drinking alcohol or caffeinated drinks^5,6,7,8^. Various comorbidities, such type 2 diabetes, and prodromal symptoms are more common among individuals who are subsequently diagnosed with PD, including anosmia, anxiety, depression, constipation, REM sleep behaviour disorder (RBD) and erectile dysfunction^7,8^. There are several examples of approaches that are being used in research settings to model the risk of PD prospectively^9–11^.

The modest overall liability explained by genetic factors and small individual effect sizes of environmental risk factors for PD suggest that interactions between them may explain some of the missing risk. Modelling interactions may yield insights into PD pathobiology, further improve prediction algorithms, and suggest potential ways to modify risk through intervention in genetically-stratified groups^12–14^.

In this study, we used the UK Biobank (UKB) cohort and the latest PD GWAS data to address three primary aims:

i. to systematically evaluate the association of environmental risk factors and prodromal features with a subsequent diagnosis of PD;
ii. to determine whether the magnitude and strength of these associations is modified by genetic risk of PD;
iii. to assess whether the inclusion of a genetic risk score improves prediction of subsequent PD compared to existing risk prediction algorithms.

## Methods

### Data sources and study design

The UKB is a large repository that contains health related data on over 500,000 individuals across the UK. The methods by which this data was collected has been described elsewhere^15^. Briefly, between 2006 and 2010 adults aged between 40 and 69 within close proximity to one of 22 UKB recruitment centres were invited to participate. Individuals had extensive demographic, lifestyle, clinical and radiological information collected. In addition to this, participants underwent genotyping and had health records collected using linked Hospital Episode Statistics.

### Study design, definition of exposures and outcomes

Our study utilised the full UKB dataset. All participants are enrolled in follow-up unless they withdraw consent, in that their medical records (Hospital Episode Statistics records, cancer register, death register, and GP records for a subset) are automatically linked to the dataset.

For analyses assessing the association of environmental and prodromal factors, we included only *incident* cases of PD (those individuals in whom the diagnosis was *recorded after* their initial assessment visit), and excluded *prevalent* cases (individuals diagnosed with PD *prior to* their baseline visit). Cases were defined as having PD if they had any record of a PD diagnosis after baseline, this was derived from self-report or linked Hospital Episode Statistics ICD codes (Supplementary Table 1). Rather than matching controls, we included all participants in the dataset as unmatched controls and adjusted for relevant confounding factors in the subsequent analyses.

All exposures were captured at the time of the initial visit. Details of how each exposure variable was defined are provided in supplementary table 1. Exposures were excluded from this analysis if the reported prevalence in UKB was substantially lower than reported population prevalence (i.e. anosmia, erectile dysfunction, shoulder pain/stiffness)^16–20^ and therefore deemed unreliably recorded. Medication-related exposures were not explored in this study because UKB does not currently have extensive linked medication data on participants.

### Demographics of cases and controls

Phenotype data were available for 2127 individuals with PD, of whom 1276 were diagnosed after enrolment (incident cases), and 500,406 controls. Of the 2127 individuals with PD, 1342 remained after exclusion of individuals of non-European ancestry and related individuals. Of the 1276 incident PD cases, 801 remained after exclusion of non-caucasian and related individuals. Of the 1276 incident cases, 1243 (97.4%) had a Hospital Episode Statistics coded diagnosis of PD, and 33 (2.59%) individuals had a self-reported diagnosis only. Demographic characteristics of individuals with PD (both prevalent and incident cases) and controls are shown in supplementary table 2. Individuals with PD were more likely to be older (mean age at recruitment 62.7 years, SD 5.49), male (61.6% male), born in the UK, of white ethnicity, less deprived, and spent longer in full-time education compared to the control group. Age at completion of full-time education was only available for a subset of participants (n=330,240). As deprivation status (which is available for all participants) is a useful proxy for socio-economic status, we chose to control for deprivation in our models to prevent exclusion of the ∼170,000 individuals with missing education data. Age at PD diagnosis was consistent with published estimates (median 66.1 years, IQR 59.5 - 71.7)^21^. Median follow-up time was the same for cases (Median 12.01, IQR 11.01 to 13.01) and controls (Median 12.01, IQR 11.01 to 13.00).

### Genotype data

Genotyping was performed using Axiom (UK Biobank Axiom™ Array, ThermoFisher) and UK BiLEVE arrays. Genotyping, imputation and quality control procedures are described elsewhere^22^. Genetic principal components were included in the UKB database (data-field 22009).

### Construction of a polygenic risk score (PRS)

A variety of PRS were created using the clumping-and-thresholding approach:

1. We extracted variant associations with PD from the most recent GWAS but not including the UKB participants from that GWAS^23^.
2. We excluded palindromic variants and variants without an rsID.
3. We excluded variants associated with PD above an arbitrary p value threshold (0.00005, 0.0005, 0.005, 0.05, 0.1, 0.2, 0.4, 0.6, 0.8, and 1).
4. We clumped using several r^2^ thresholds (0.1, 0.2, 0.4, 0.6, 0.8) and a clumping distance of 250kb, with the 1000 genomes EUR samples as the reference genome^24^.

Reference genome data were obtained from the 503 participants of European ancestry in the 1000 genomes project^25^. Only autosomal, biallelic variants which passed quality control in both the PD GWAS and target (UKB) datasets were included. We excluded all duplicate rsIDs, duplicate positions, variants deviating from Hardy-Weinberg Equilibrium (p <1e-06), rare variants with minor allele frequencies <0.01, variants with genotype missingness >10%, and variants with low imputation quality (Mach R^2^ < 0.3). After SNP QC, a total of 4,490,455 markers overlapped between the reference and target datasets. For genetic analysis, individuals with >10% missing genotypes were excluded, and only individuals with self-reported ‘British’ ethnicity and genetically European ancestry as defined by genetic principal components were included. We excluded one of each pair of individuals related at a kinship coefficient cutoff of 0.0442, equivalent to a 3rd degree relative, e.g. first cousins. Kinship coefficients were calculated by UKB and are provided in the ‘Relatedness’ file available for download (category 263).

As a sensitivity analysis to determine whether as-yet-undiscovered genomic risk loci explained additional liability to PD, we created an additional PRS using the best-performing PRS (in terms of Nagelkerke’s pseudo-R^2^). We excluded all variants within 1MB either side of the lead SNP for the 90 risk loci discovered in the most recent IPDGC GWAS ^4^.

Effect allele dosage at each locus was multiplied by the beta coefficient to generate the risk score for that locus. Scores were standardised to have mean 0 and unit variance for each SNP. For missing genotypes, the score at that locus was defined as the mean of all scores at that locus. Risk scores were totalled across the genome to calculate an individual’s score. All individuals in the UKB with a PD diagnosis, prevalent or incident, were included. Analysis was performed in PLINK (v2.00aLM 64-bit Intel) using the ‘--score’ flag.

### Statistical methods

Multivariate models were built for each risk factor using the entire UKB cohort as controls and adjusting for age, sex, ethnicity, and current deprivation status. Secondly, a multivariate logistic regression model for incident PD comprising all environmental factors with robust associations to PD risk was built, including the above confounders. Likelihood ratio tests were used to assess the improvement of model fit at a False Discovery Rate threshold of 0.05.

Interactions were assessed on both the additive and multiplicative scales. Interaction on the additive scale was assessed by calculating the Attributable Proportion due to interaction (AP). Additive interaction analyses were based on multivariate logistic regression models incorporating age at recruitment, sex, and the first four genetic principal components as confounders^26^.

For a logistic regression model of the form: 

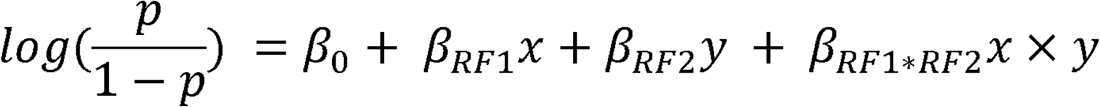

In which 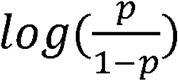 is the log odds of PD, *x* and *y* are the values of exposure variables (e.g. childhood body size, smoking, polygenic risk score), and *x* × *y* is the interaction term, then the Relative Excess Risk due to Interaction (RERI) can be calculated as: 

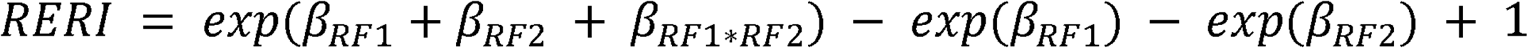

The AP can be conceived of as the proportion of the disease in the doubly-exposed group attributable to the interaction between the risk factors, i.e: 

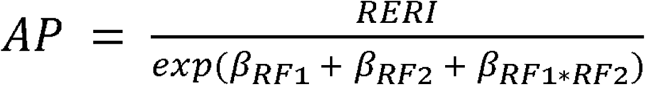

This model can be expanded to include confounding covariates, in which case the beta coefficients are adjusted for confounders. Derivation and further discussion of the advantages of this method over Rothman’s initial description can be found in Knol et al^26^. We restricted this analysis to participants with genetically European ancestry determined by both self-report (“Caucasian” in UKB data) and genetic ethnic grouping. For interaction analyses using the PRS, covariates were age, sex, current deprivation, and the first four genetic principal components. The PRS was transformed using the inverse-normal transformation and treated as a continuous variable for these analyses. For the menarche analysis, age at menarche was also transformed using the inverse normal transformation. Confidence intervals for the AP were estimated using bootstrap resampling of the entire dataset with replacement for 5000 iterations^26^. 95% confidence intervals were derived from the 2.5th and 97.5th percentile values. Interaction on the multiplicative scale was assessed using a logistic regression model incorporating an interaction term. The presence of multiplicative interaction was assessed using the likelihood ratio test.

### Application of the PREDICT-PD algorithm

We applied the PREDICT-PD algorithm to UKB participants to externally validate this risk score and determine whether its predictive performance was enhanced by the addition of a genetic risk score^10^. Baseline risk of PD (on the odds scale) was determined from the following equation^10^: 

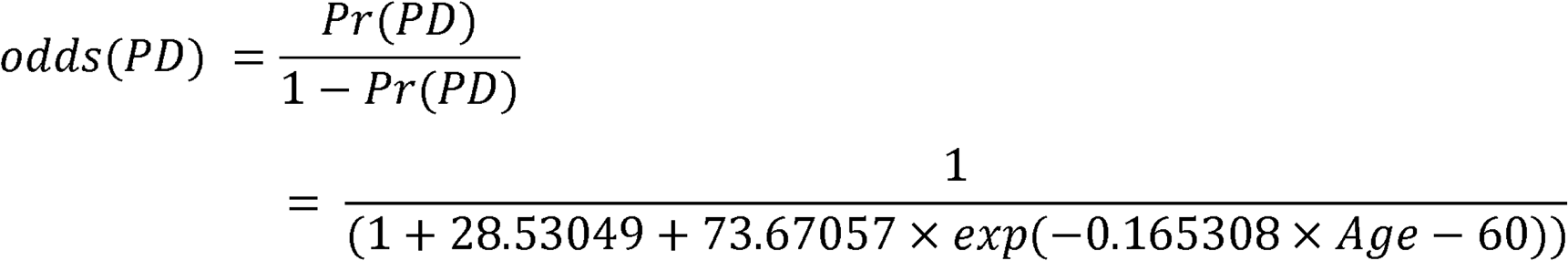

With the PREDICT-PD algorithm, the following adjustments to this baseline age-adjusted risk are made for individuals based on the presence or absence of the following traits^10^: females (divided by 1.5), current smoking (multiplied by 0.44), previous smoking (multiplied by 0.78), family history of PD (multiplied by 4.45), more than one cup of coffee per day (multiplied by 0.67), more than one alcoholic drink per week (multiplied by 0.9), constipation (multiplied by 2.34), anxiety or depression (multiplied by 1.86), and erectile dysfunction (multiplied by 3.8). The final odds for PD was converted to the probability scale using the equation: 

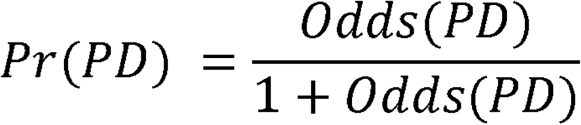

### Ethical approval

This work was performed using data from UKB (application number 14872). All participants gave informed consent to UKB registration and are free to withdraw from the study at any point, at which point their data are censored and cannot be included in further analyses.

### Computing

This research was supported by the High-Performance Cluster computing network hosted by Queen Mary University of London^27^.

Statistical analyses were performed in R version 3.6.1 using RStudio version 1.2.1335. Extraction of European individuals from the 1000 genomes reference genome was conducted using vcftools. Construction of the polygenic risk score, application of the polygenic risk score to individuals, and quality control were performed in PLINK v1.9 and PLINK v2.00.

## Results

### Risk factors and prodromes

There was strong evidence of a positive association (P_adjusted_< 0.05; 32 tests) between incident PD diagnosis and having a family history of PD, not smoking, low alcohol consumption (<1 drink / week), depression, excessive daytime sleepiness, a family history of dementia, epilepsy and earlier menarche. There was weaker evidence (FDR < 0.10) for an association between PD and having had peptic ulcer disease or diabetes mellitus (Fig 1, Table 1). Effect estimates and precision did not alter substantially in a multivariate model including all strongly-associated (FDR < 0.05) risk factors (age of menarche was excluded to allow for inclusion of both sexes; Table 2).

**Figure 1:**
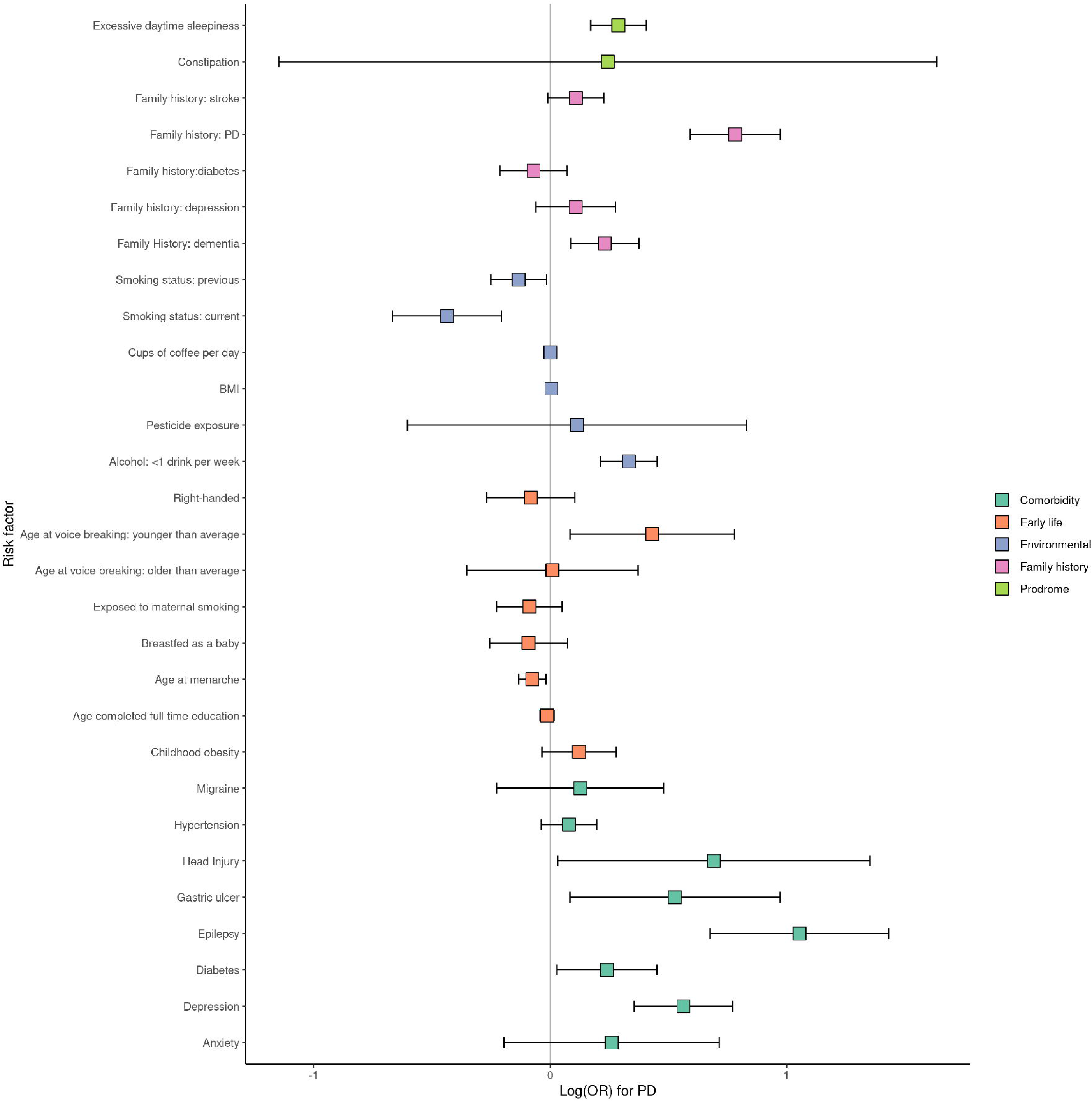
Associations of risk factors and incident cases of Parkinson’s disease. Point estimates for association are depicted as log odds ratios and 95% confidence intervals. Estimates of association were derived from logistic regression models adjusting for age, sex, Townsend deprivation index at recruitment, and ethnicity. BMI = body mass index; PD = Parkinson’s disease.

### Validation of PREDICT-PD risk algorithm

We sought to validate a risk prediction algorithm which has previously been employed in a longitudinal cohort study of UK residents to determine risk of PD (the PREDICT-PD study; www.predictpd.com)^9^. The algorithm uses published estimates of relative risks and odds ratios derived from large meta-analyses of early non-motor features and risk factors for PD^7^. In the present study, the algorithm had discriminative ability for distinguishing incident PD cases from controls (Nagelkerke’s pseudo-R^2^ 0.035, likelihood ratio *p* < 2×10^−16^). The median predicted odds of PD was 2.68x higher among incident PD cases (Median odds 0.00957, IQR 0.0110) compared to controls (Median odds 0.00357, IQR 0.00598; Fig 2)^28^.

**Figure 2:**
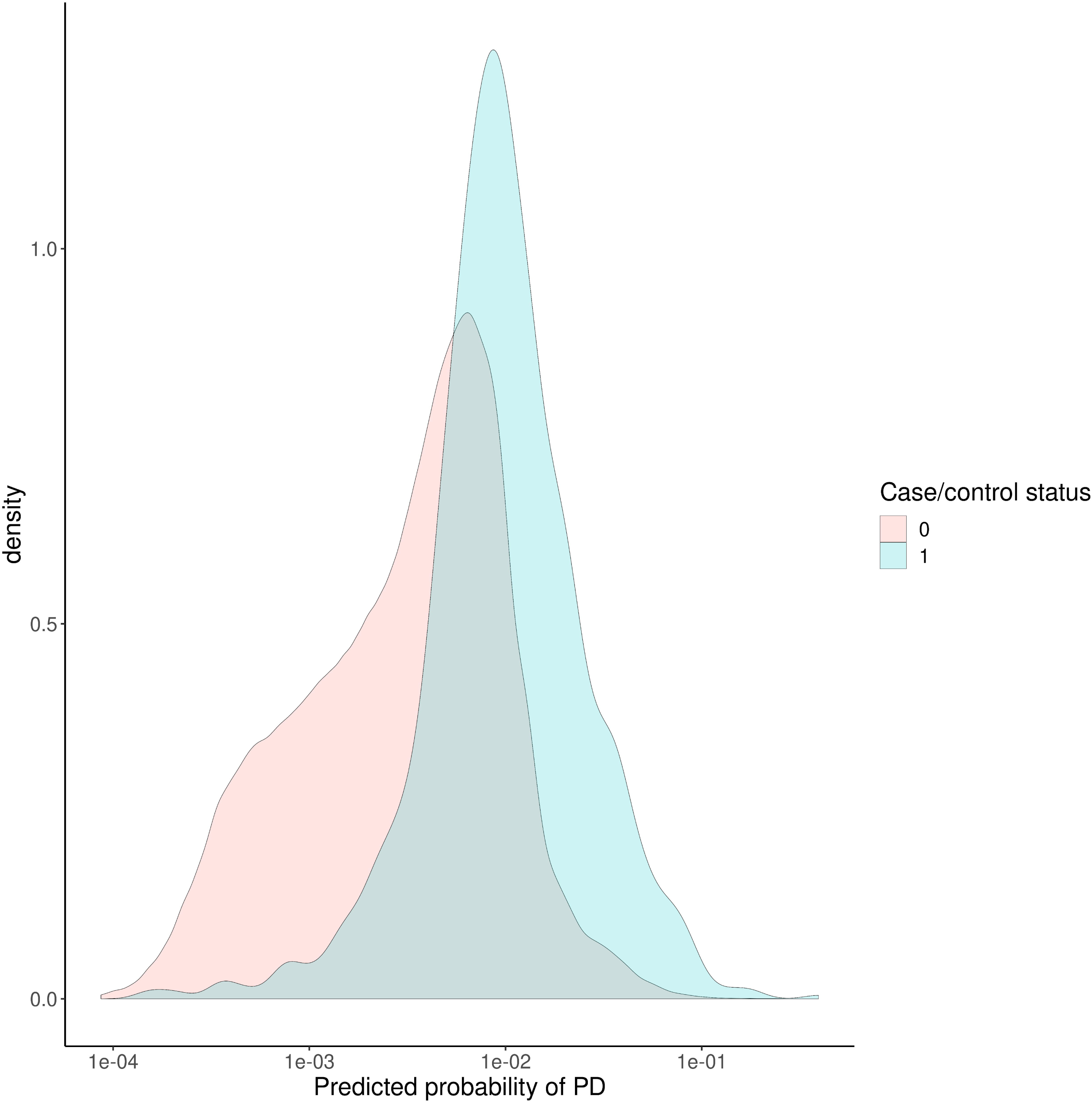
PREDICT-PD determined probability (on the absolute risk scale) of Parkinson’s disease, determined at recruitment, for individuals who would go on to develop PD (incident cases) and those who would not (controls).

### Genetic risk score

Next, we built polygenic risk scores for PD using the clumping-and-thresholding approach. We used several different P value and R^2^ thresholds to generate multiple scores, and selected the score with the closest fit to the UK Biobank data as measured by Nagelkerke’s pseudo-R^2^. The parameters of the best-fitting PRS were: p value threshold < 5×10^−4^, clumping R^2^ threshold 0.8, Nagelkerke’s pseudo-R^2^ 0.011, 4285 SNPs included.

For this PRS, individuals with the highest 10% of scores had roughly 3.3x increased risk of PD (OR 3.30, 95% CI 2.57 - 4.24) compared to the lowest risk decile. Higher PRS scores were associated with earlier age at PD diagnosis in a linear model adjusting for age, sex and the first four genetic principal components (PCs; beta −0.73 per 1-SD increase in PRS, *p* = 0.002, Fig 3) - this estimate is similar to the published estimate from the IPDGC^29^. Inclusion of the PRS improved model fit compared to a null model including only the PREDICT-PD algorithm, age, sex, the first four genetic PCs (Nagelkerke pseudo-R^2^ 0.0053, p = 6.87×10^−14^). The PRS therefore improves the performance of the PREDICT-PD algorithm, which in its current form does not include explicit genetic data (it does include family history, which is an imperfect surrogate for genetic risk). We modified this PRS to exclude all variants within known PD genomic risk loci (supplementary table 3): for each risk locus, all variants 1MB either side of the lead SNP were removed. This modified PRS also explained additional PD liability compared to the PREDICT-PD algorithm alone (Nagelkerke pseudo-R^2^ 2.57×10^−3^, p = 1.63×10^−7^), suggesting that as-yet-undiscovered PD risk loci may explain additional risk not accounted for by known loci and other risk factors incorporated in the PREDICT-PD algorithm.

**Figure 3A:**
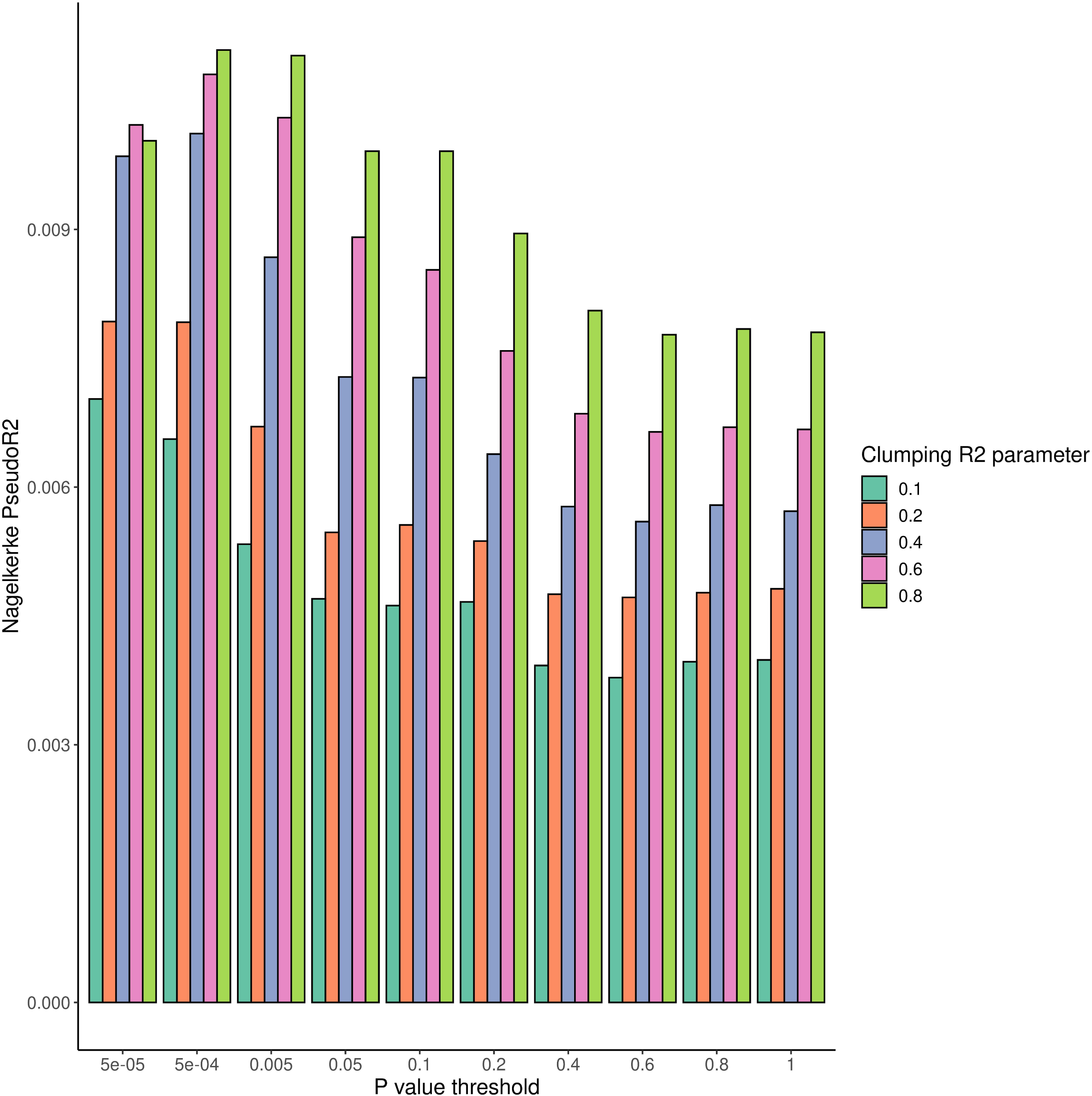
Several candidate polygenic risk scores (PRS) were created using summary statistics from the Meta5 PD GWAS excluding UKB participants. For each candidate PRS, the degree of variation in PD risk explained was estimated using Nagelkerke’s pseudo-R2 metric. B: Normalised PRS values for incident PD cases and controls. C: Odds ratio of Parkinson’s disease by polygenic risk score decile compared to lowest polygenic risk score decile. D: correlation between increasing PRS and earlier age at PD diagnosis. E: Interactions between risk factors for PD and the PD PRS were estimated using the Attributable proportion due to interaction. Point estimates for the AP and 95% CIs are shown. OR = odds ratio; PD = Parkinson’s disease; PRS = polygenic risk score

### Interactions

We looked for interactions between the genome-wide PRS and the 10 risk factors/prodromal symptoms found to be associated with PD risk at FDR < 0.10. The only phenotype with strong evidence (FDR P < 0.05) for a negative multiplicative interaction was between diabetes and the PRS (beta −0.47, FDR P = 0.004, supplementary table 4) - i.e. diabetes appears to increase PD risk for individuals with a low PRS, but has a substantially weaker effect - and may even be protective - for individuals with a high PRS. We also only found strong evidence of interaction on the additive scale between the PRS and diabetes (AP −0.42, 95% CI −0.81 to −0.11; Fig 3). These results suggest that diabetes is a more potent risk factor among people at low genetic risk of PD.

To illustrate the interaction between the PRS and diabetes, we built regression models (PD status ∼ Age + Sex + 4 genetic PCs + Diabetes) using participants in the highest and lowest deciles for the PRS (i.e. the bottom and top 10% of prior genetic risk). Among the high genetic risk group, the effect of diabetes was imprecisely estimated but suggested a protective effect (OR = 0.28, 95% CI 0.07 - 1.15, p = 0.078), whereas among the low genetic risk group the effect was opposite (OR = 2.76, 95% CI 1.22 - 6.27, p = 0.015).

## Discussion

Here, we used publicly-available data from the UKB cohort study to examine risk and protective factors for PD. It is widely believed that the earlier in the pathological course of PD that a disease-modifying intervention will be used, the better the chance of delaying symptom onset or preventing phenoconversion.

We observed strong associations with incident PD for several well-established risk and protective factors. In a model adjusted for age, sex, ethnicity and current deprivation, risk factors for incident PD where: having a family history of PD, not smoking, low alcohol consumption (<1 drink / week), depression, excessive daytime sleepiness, a family history of dementia, epilepsy, and earlier age of menarche. All of these factors remained associated with incident PD when modelled with all other potential exposures. Previously reported associations that fell short of the FDR Q value were preceding gastric ulcer or diabetes mellitus^7^. Notable exposures for which there was no evidence of effect were anxiety, BMI, constipation, pesticide exposure and coffee consumption.

In a recent paper from some members of this group, ‘novel’ cross-sectional associations with PD were reported for migraine and epilepsy^8^. Here we have not only replicated the association with epilepsy, but we have also demonstrated a temporal relationship with incident PD. Whether the association is driven by epilepsy or chronic use of anti-epileptic drugs remains to be determined. As linked medication data becomes available, we will be able to clarify the drivers of this association. There was no convincing association with migraine in the present study.

The association of earlier age at menarche with PD is novel and intriguing. In addition, we observed weak evidence for a similar effect of earlier age at voice breaking in males, suggesting that earlier pubertal timing in both sexes may increase PD risk. The sex dimorphism in PD incidence suggests possible protective roles for female sex hormones, or possible harmful roles for male sex hormones^30^. In animal models of PD, sex hormones have pleiotropic effects which are inconsistent between studies^30–34^. The broad consensus from animal models and epidemiological studies of menopausal timing and PD^35,36^ is that oestrogens may be neuroprotective. In this context, our findings are counterintuitive, as earlier menarche should predispose towards greater lifetime oestrogen exposure. It is possible that the observational association between earlier puberty and PD risk is driven by residual confounding. Both the genetic and environmental determinants of pubertal timing may confound the relationship with PD risk^37^. Thus we would interpret this association with caution and encourage replication of this finding in other cohorts.

Next, we demonstrated that a basic risk algorithm previously-developed in the PREDICT-PD study could be used to identify incident cases of PD in UKB. In separate work, we have observed that whilst the basic PREDICT-PD algorithm has utility, performance significantly improves when factors such as anosmia, probable RBD and subtle motor impairment are modeled as part of the algorithm (*unpublished data*). UKB does not accurately capture data on these three exposures, so we were not able to include them. However, in the current study, we were able to extend the prediction concept to show that the addition of genetic liability towards PD (in the form of a PRS) improved the performance of the basic algorithm. Whilst individual risk estimates may not be particularly meaningful, this approach can be used to define a higher risk group for testing disease-modifying strategies^38^. The use of a PRS with and without other risk factors for PD has been previously validated in a large case-control setting^39^, but there are limited examples of application in a population setting such as we have done ^40^. We anticipate that in a prospective setting in which there is information for the basic algorithm, data on smell, RBD and motor dysfunction, and polygenic risk, then performance may increase considerably.

Finally, we undertook some preliminary study of the role of gene-environment interactions for PD in UKB. We compared how the association of the various exposures in the model varied across deciles of genetic risk. Prior to this, simple gene-environment interaction studies have been undertaken to investigate the effect modification that a change at a single gene or locus can have on an environmental risk factor^41,42^. Here, we used the aforementioned PRS to show that the association with diabetes is potentially modified such that it plays a bigger role as a risk factor in those at lower genetic risk and may have (or its treatment may have) a protective effect in those at higher genetic risk.

This observation is especially interesting in the context of recent phase II clinical trial data showing that the anti-hyperglycaemic drug exenatide (a Glucagon Like Peptide 1 agonist) had efficacy in reducing off-medication motor symptoms in PD^43^. It is conceivable that our results may therefore reflect confounding by drug treatment - i.e. if the treatment of diabetes differs systematically between individuals at high and low risk of PD. As genetic risk for PD (quantified by genome-wide PRS) may itself be a surrogate for subtle ethnic variation, socio-economic status and other confounders, so it is plausible that there could be real differences in access to particular anti-diabetes medications between strata of the PRS. If the effect of anti-diabetic drugs on PD is modified dramatically by prior genetic risk for PD, it may be possible to select individuals who are more likely to benefit from these drugs in phase III trials. Validation of our results and exploration of the mechanism for this interaction is required before translation into trial selection criteria.

We have previously observed markedly different effects of diabetes on PD risk^7^. Whilst survival bias may account for some of the observed variability in effect estimates comparing case-control and cohort studies, genetic population stratification may also be an important source of variation as indicated here. Correcting for genetic principal components should mitigate confounding due to population stratification, but may not eliminate it. The importance of genetic stratification for PD intervention studies, has been recently explored^38^, and in the current study, we demonstrate further evidence for why genetic stratification is an important consideration.

The strengths of this study are that we used a very large sample size to measure risk and protective factors for PD, as well as to externally validate the PREDICT-PD algorithm in a cohort where incident cases are accruing. The prospective design reduces the likelihood of reverse causation but in diseases with a long prodromal phase (such as PD), reverse causation cannot be completely dispelled. However, for the purpose of predicting incident cases, whether factors in the model are true exposures or prodromal features is less concerning. As many of the exposures vary by age and gender, we adjusted all exposure variables for important confounding factors. We have previously surveyed a sub-group of the PREDICT-PD participants and found that less than 5% were participants in UKB, hence overlap in populations is minimal. The latest PD GWAS used data from PD cases in UKB and the controls, but we used summary statistics which excluded UKB cases, controls and proxy-cases, to avoid sample overlap and over-fitting models.

General limitations are that definition of incident PD cases in this setting relied to a small extent on self-report and several important risk factors for incident PD were inadequately captured. Both of these factors may lead to bias and imprecision in the effect estimates. Another important consideration is the generalisability of UKB. Recruitment into the UKB cohort was voluntary with 5.5% of those invited ultimately joining. Comparing the UKB population to UK Census and representative cross-sectional survey data shows that typically UKB participants were more likely to be female, older and from less socioeconomically deprived areas, within the cohort rates of smoking, obesity and daily drinking were less than that in the general UK population^44^.

To conclude, we have further confirmed several well-established risk and protective factors for PD, and shed further light on several novel associations (migraine, epilepsy, earlier menarche). We have externally validated the basic PREDICT-PD algorithm and extended this approach to incorporate population-level common genetic variation. Finally we have modelled interactions between environmental factors, comorbidities and polygenic risk to demonstrate further improvement in model fit and conceptually how genetic stratification might aid or confound intervention strategies.

## Acknowledgements

This research has been conducted using the UK Biobank Resource under Application Number 14872.

## Author Contributions

B.M.J, D.B, A.J.N designed the study methods and wrote the first draft of the manuscript. J.P.B, C.B, S.BC, K.H, R.D, M.A.N, A.B.S, A.S reviewed the study methods and statistical analysis. R.D, J.H, G.G, A.J.L, A.S contributed in the discussion and reviewed and modified the manuscript.

---

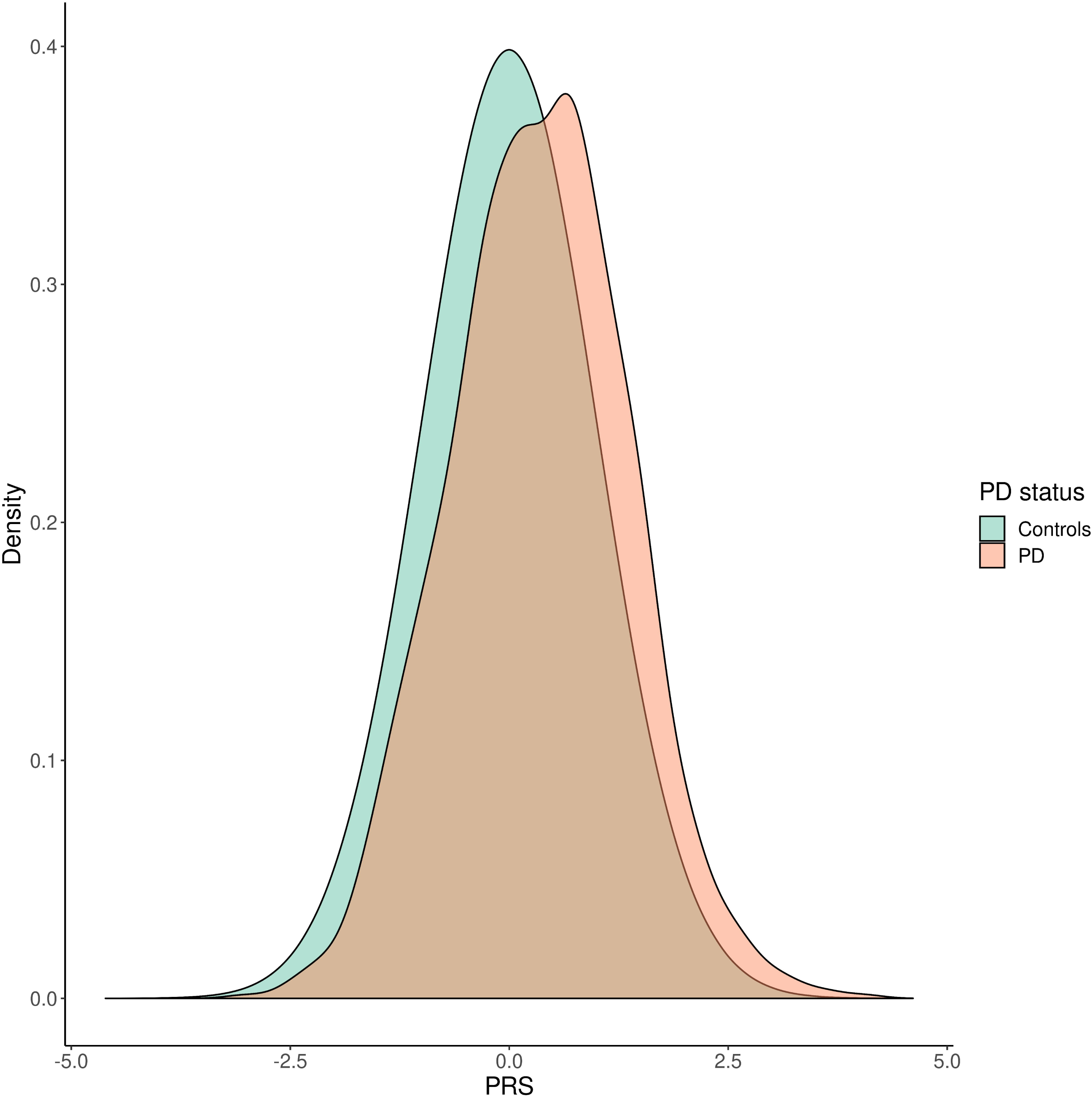

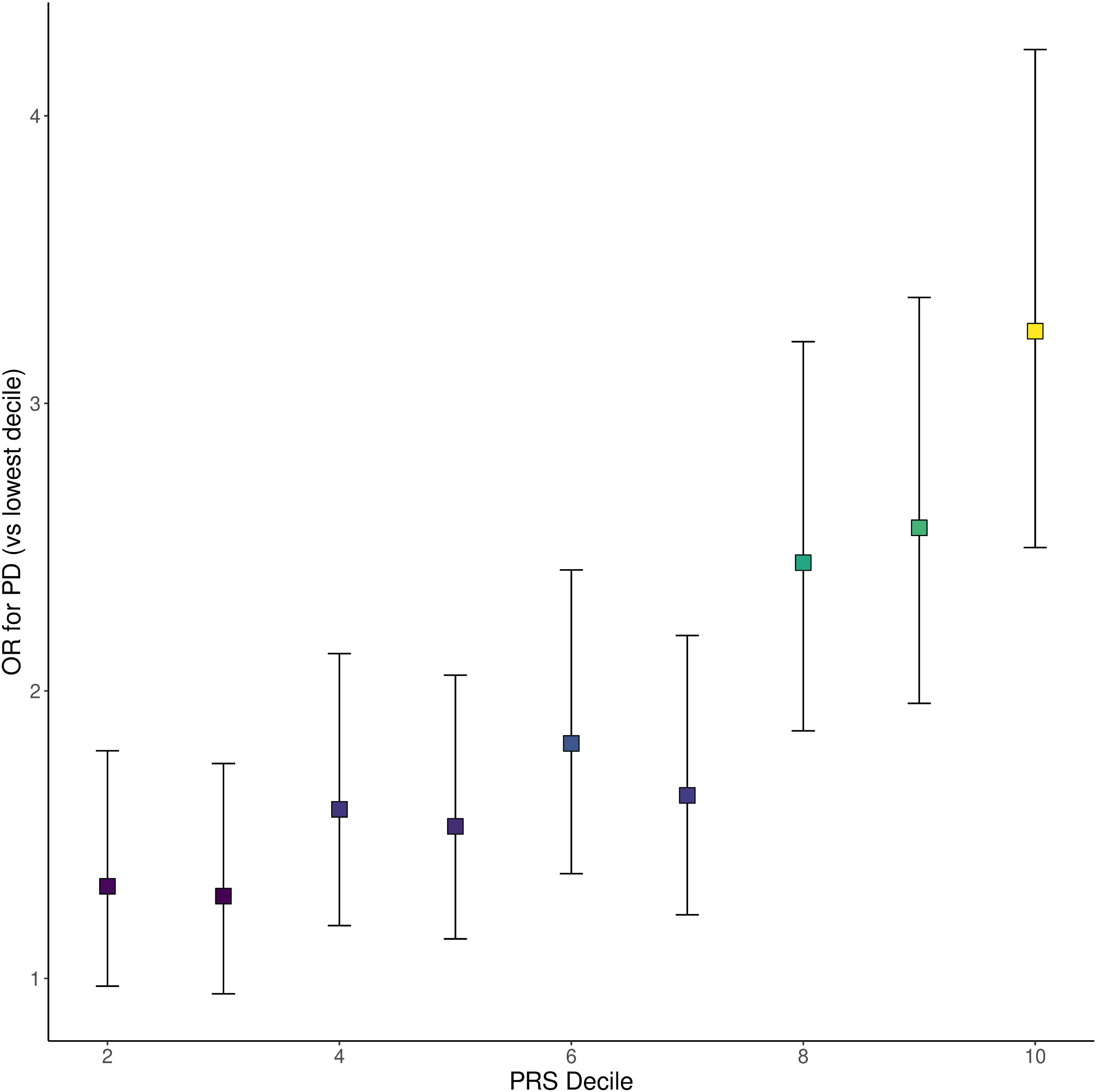

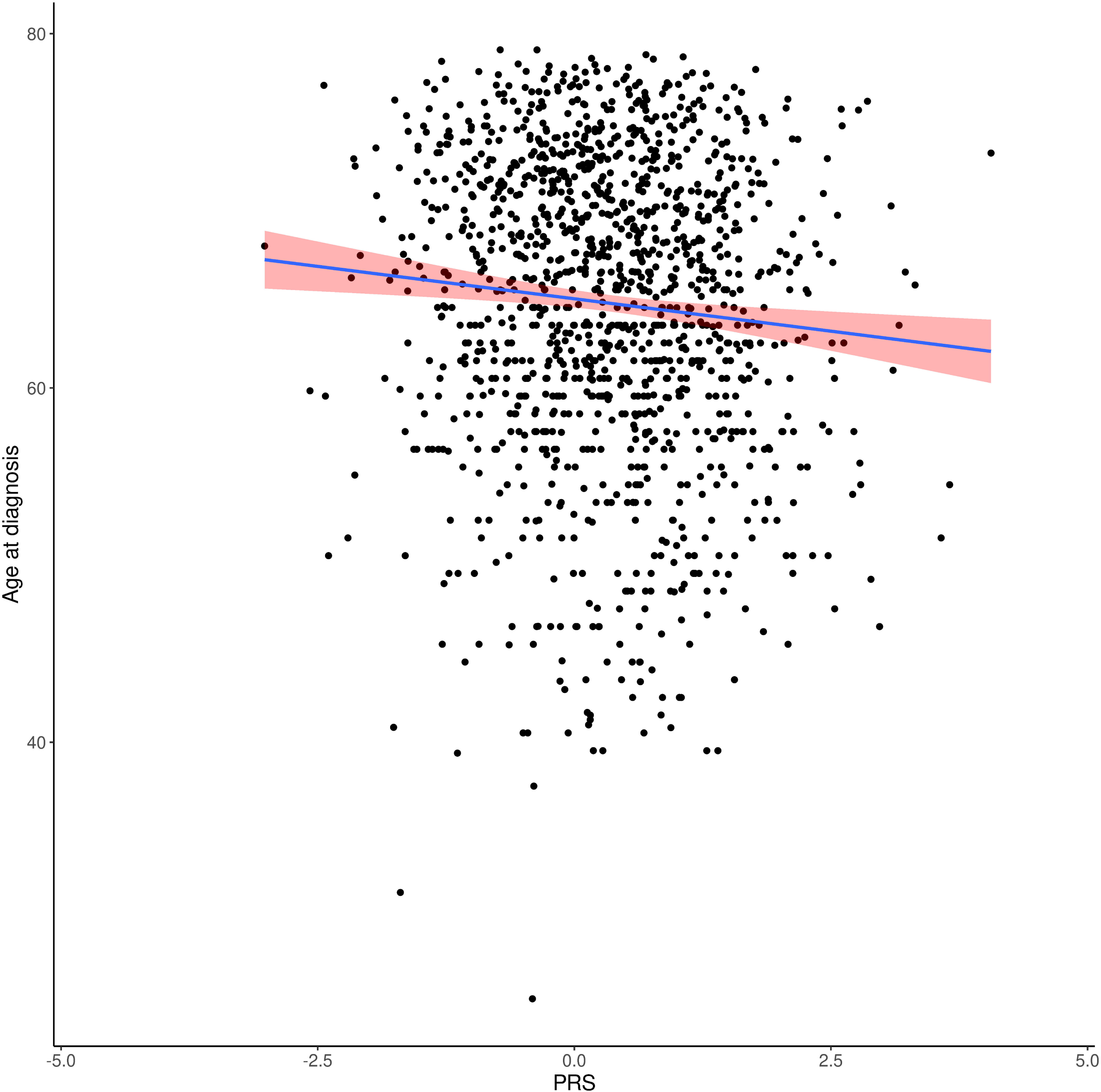

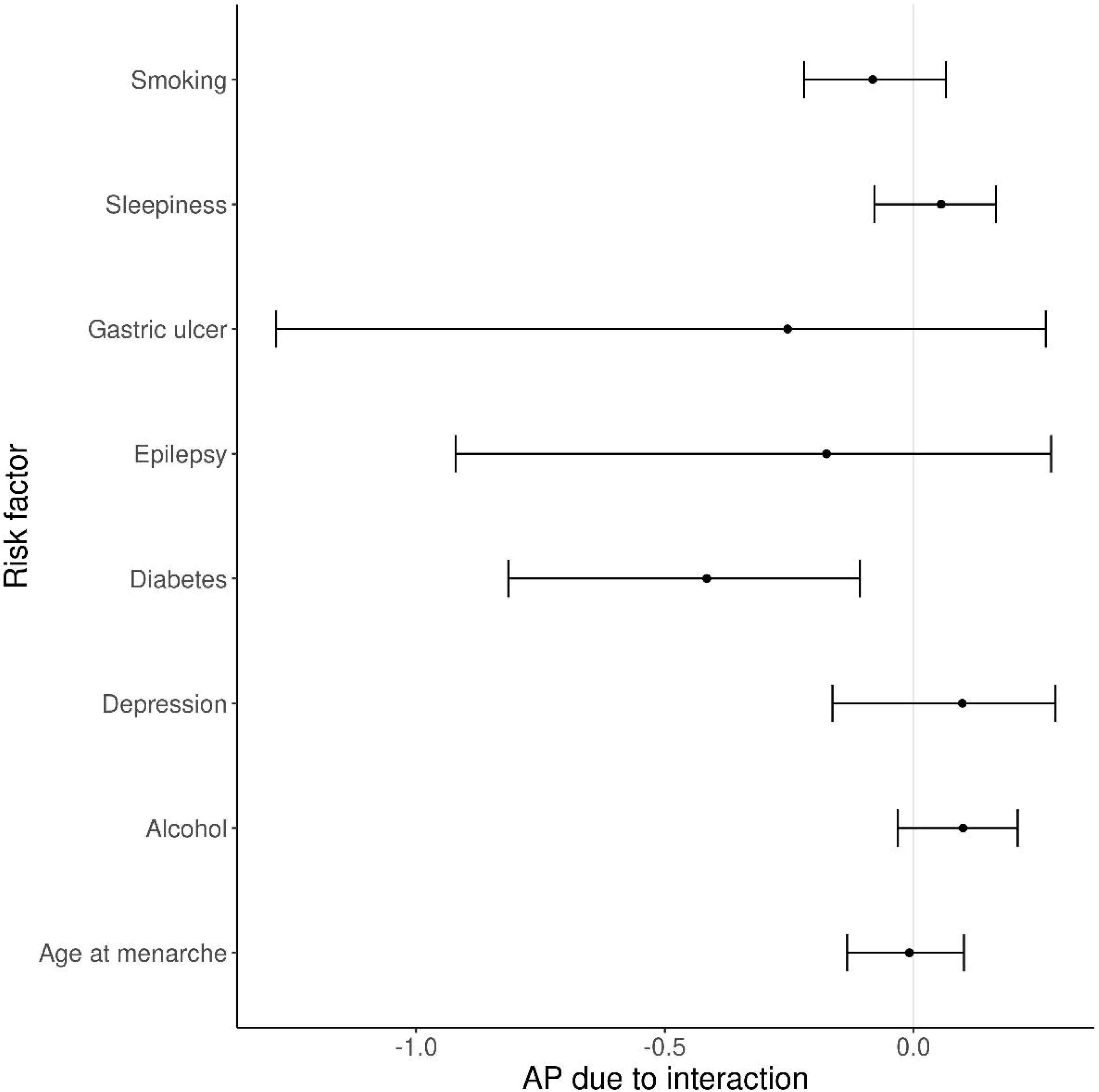

